# *In vivo* antiviral host response to SARS-CoV-2 by viral load, sex, and age

**DOI:** 10.1101/2020.06.22.165225

**Authors:** Nicole A. P. Lieberman, Vikas Peddu, Hong Xie, Lasata Shrestha, Meei-Li Huang, Megan C. Mears, Maria N. Cajimat, Dennis A. Bente, Pei-Yong Shi, Francesca Bovier, Pavitra Roychoudhury, Keith R. Jerome, Anne Moscona, Matteo Porotto, Alexander L. Greninger

**Affiliations:** Department of Laboratory Medicine, University of Washington School of Medicine, Seattle, WA, USA; Galveston National Laboratory, University of Texas Medical Branch, Galveston, Texas, 77550; Department of Experimental Pathology, University of Texas Medical Branch, Galveston, Texas, 77550; Department of Microbiology and Immunology, University of Texas Medical Branch, Galveston, Texas, 77550; Department of Biochemistry and Molecular Biology, University of Texas Medical Branch, Galveston, Texas, 77550; Center for Host–Pathogen Interaction, Columbia University Medical Center, New York, New York, 10032, USA; Vaccine and Infectious Disease Division, Fred Hutchinson Cancer Research Center, Seattle, WA; Department of Pediatrics, Columbia University Medical Center, New York, New York, USA; Department of Microbiology & Immunology, Columbia University Medical Center, New York, New York, 10032, USA; Department of Physiology & Cellular Biophysics, Columbia University Medical Center, New York, New York, 10032, USA; Department of Experimental Medicine, University of Campania “Luigi Vanvitelli”, 81100 Caserta, Italy

**Author notes:** Corresponding Author **Correspondence**: Alexander L. Greninger, 1616 Eastlake Ave E, Suite 320, Seattle, WA 98102.

**Keywords:** SARS-CoV-2, COVID-19, RNAseq, interferon, immune response, ribosomal proteins, *ACE2*

## Abstract

Despite limited genomic diversity, SARS-CoV-2 has shown a wide range of clinical manifestations in different patient populations. The mechanisms behind these host differences are still unclear. Here, we examined host response gene expression across infection status, viral load, age, and sex among shotgun RNA-sequencing profiles of nasopharyngeal swabs from 430 individuals with PCR-confirmed SARS-CoV-2 and 54 negative controls. SARS-CoV-2 induced a strong antiviral response with upregulation of antiviral factors such as *OAS1-3 and IFIT1-3*, and Th1 chemokines *CXCL9/10/11*, as well as a reduction in transcription of ribosomal proteins. SARS-CoV-2 culture in human airway epithelial cultures replicated the *in vivo* antiviral host response. Patient-matched longitudinal specimens (mean elapsed time = 6.3 days) demonstrated reduction in interferon-induced transcription, recovery of transcription of ribosomal proteins, and initiation of wound healing and humoral immune responses. Expression of interferon-responsive genes, including *ACE2*, increased as a function of viral load, while transcripts for B cell-specific proteins and neutrophil chemokines were elevated in patients with lower viral load. Older individuals had reduced expression of Th1 chemokines *CXCL9/10/11* and their cognate receptor, *CXCR3*, as well as CD8A and granzyme B, suggesting deficiencies in trafficking and/or function of cytotoxic T cells and natural killer (NK) cells. Relative to females, males had reduced B and NK cell-specific transcripts and an increase in inhibitors of NF-κB signaling, possibly inappropriately throttling antiviral responses. Collectively, our data demonstrate that host responses to SARS-CoV-2 are dependent on viral load and infection time course, with observed differences due to age and sex that may contribute to disease severity.

## Introduction

The novel coronavirus SARS-CoV-2 that emerged in late 2019 from Wuhan, China, has rapidly spread throughout the world, causing more than 6 million cases and 400,000 deaths globally as of June 2020. COVID-19 morbidity and mortality has been overwhelmingly concentrated in elderly individuals and those with preexisting comorbidities (1). In older individuals, immunosenescence and dysregulated antiviral responses due to viral chronic low-grade age-related inflammation may play an important role (2), as has been proposed for influenza (3). Males are known to be generally more susceptible to infectious disease than females (4) and SARS-CoV-infected male mice had increased infiltration of inflammatory macrophages into their lungs, leading to a deleterious inflammatory response (5). Accordingly, systemic inflammatory markers such as neutrophil-to-lymphocyte ratio and C-reactive protein were elevated in men who died of SARS-CoV-2 (6). However, the mechanisms behind increased mortality among older adults and males with COVID-19 remain speculative.

Entry of SARS-CoV-2 into host cells depends on binding to the receptor ACE2 (7), expressed at a high level in the nasal epithelium (8), then further induced upon exposure to interferon (9), suggesting a mechanism by which SARS-CoV-2 exploits host antiviral responses. SARS-CoV antagonizes initial viral detection and interferon responses by an as-yet unknown mechanism (10,11). SARS-CoV-2 may employ similar mechanisms, as low MOI infections of bronchial epithelial cells do not result in extensive transcription of interferon-stimulated genes (ISGs) at 24 hours post infection (12). An important consequence of these observations is that SARS-CoV-2 viral load and transmissibility peaks at the time of symptom onset (13,14). The temporal relationship between viral load and host gene expression has not been fully explored.

In the United States, diagnostic testing is generally performed on nasopharyngeal (NP) swabs, from which SARS-CoV-2 RNA can be recovered. Shotgun RNA sequencing of this material allows for simultaneous recovery of viral genomes for transmission tracking as well as understanding of *in situ* host response (15). Since the first detection of SARS-CoV-2 in the USA in WA State, the University of Washington Virology Laboratory has performed shotgun RNA sequencing to recover more than 1,000 viral genomes to understand the evolution and molecular epidemiology of the virus (16,17). Here, we examine host specific gene expression differences by SARS-CoV-2 infection status, host age, sex, and viral load in nasopharyngeal swabs from 430 SARS-CoV-2 infected individuals and 54 negative controls.

## Results

Since early March, 2020, the University of Washington Virology Laboratory has tested more than 100,000 samples, primarily NP swabs, for infection with SARS-CoV-2 (18). Thousands of SARS-CoV-2 positive samples, as well as negative controls, have subsequently been metagenomically sequenced, contributing to a detailed understanding of the phylogeny and molecular epidemiology of the virus (16,19). In this study, we selected a subset of sequenced samples that had sufficient reads (>500,000) pseudoaligned to the human transcriptome to examine gene expression changes as a result of RT-PCR-confirmed SARS-CoV-2 infection. Patient demographics are summarized in Table 1.

**Table 1.**
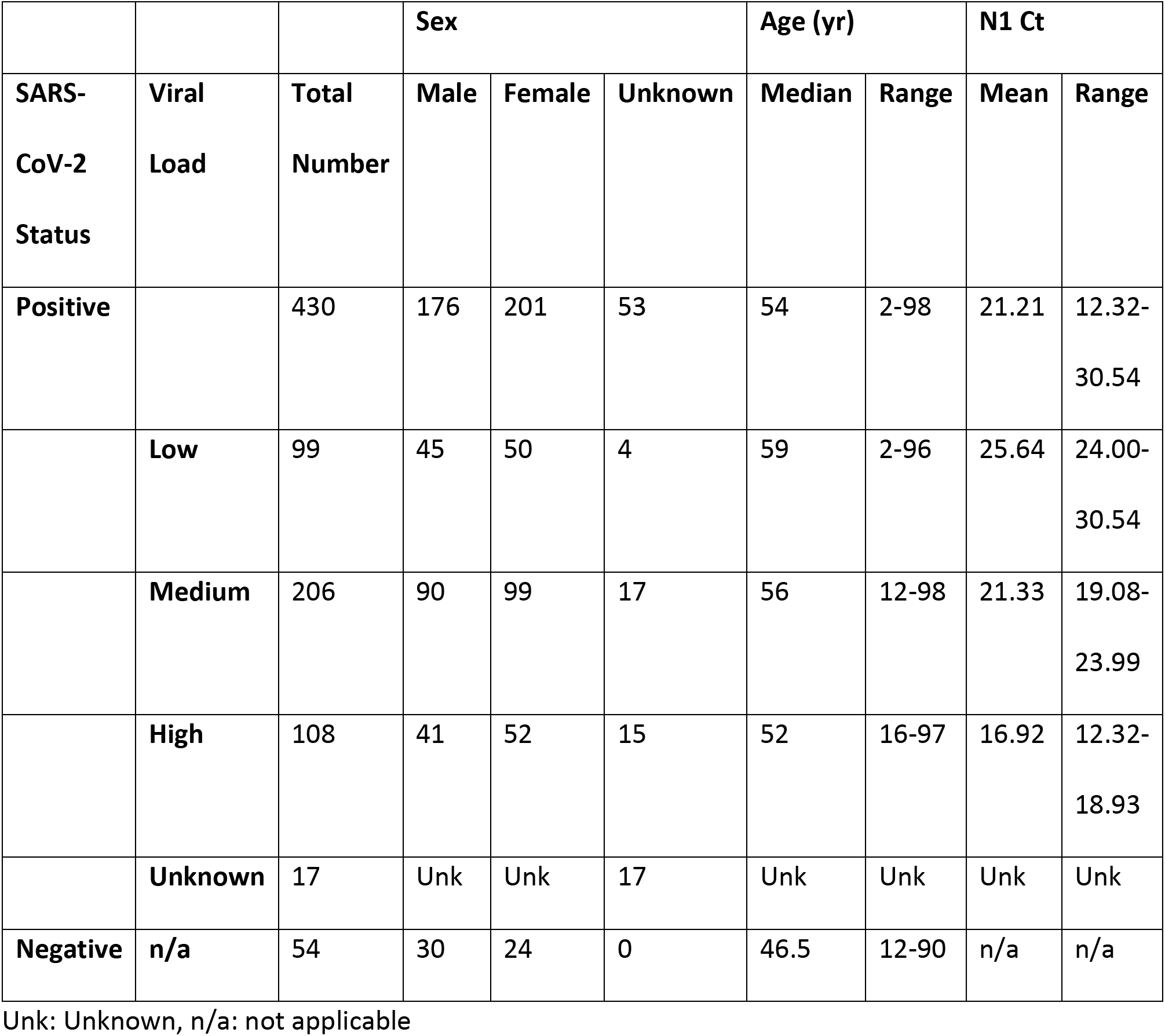
Patient demographics of SARS-CoV-2 positive and negative samples.

|  |  |  | Sex |  |  | Age (yr) |  | N1 Ct |  |
| --- | --- | --- | --- | --- | --- | --- | --- | --- | --- |
| SARS-CoV-2 Status | Viral Load | Total Number | Male | Female | Unknown | Median | Range | Mean | Range |
| Positive |  | 430 | 176 | 201 | 53 | 54 | 2-98 | 21.21 | 12.32-30.54 |
|  | Low | 99 | 45 | 50 | 4 | 59 | 2-96 | 25.64 | 24.00-30.54 |
|  | Medium | 206 | 90 | 99 | 17 | 56 | 12-98 | 21.33 | 19.08-23.99 |
|  | High | 108 | 41 | 52 | 15 | 52 | 16-97 | 16.92 | 12.32-18.93 |
|  | Unknown | 17 | Unk | Unk | 17 | Unk | Unk | Unk | Unk |
| Negative | n/a | 54 | 30 | 24 | 0 | 46.5 | 12-90 | n/a | n/a |

Unk: Unknown, n/a: not applicable

We first characterized the genes most differentially expressed (DE) in the nasopharynx as a result of SARS-CoV-2 infection (n=430 positive, 54 negative). After correcting for batch effects, we found 83 differentially expressed genes (padj <0.1 and absolute log2FoldChange >1) between SARS-CoV-2 positive and negative samples, comprising 41 upregulated genes and 42 downregulated genes (Supplementary Table 1). Clustering of samples by the 50 most significant DE genes reveals multiple gene expression clusters among SARS-CoV-2 positive samples, while most negative samples cluster together (Figure 1A). Consistent with results from Butler et al (20), SARS-CoV-2 infection induces an interferon-driven antiviral response in the nasopharynx, upregulating transcripts encoding viral sensors (*DDX60L*), chemokines that attract effector T cells and NK cells (*CXCL9*, *10*, *11*), and direct inhibitors of viral replication and function (*MX2*, *RSAD2*, *HERC5*), highlighted in Figure 1B

**Figure 1:**
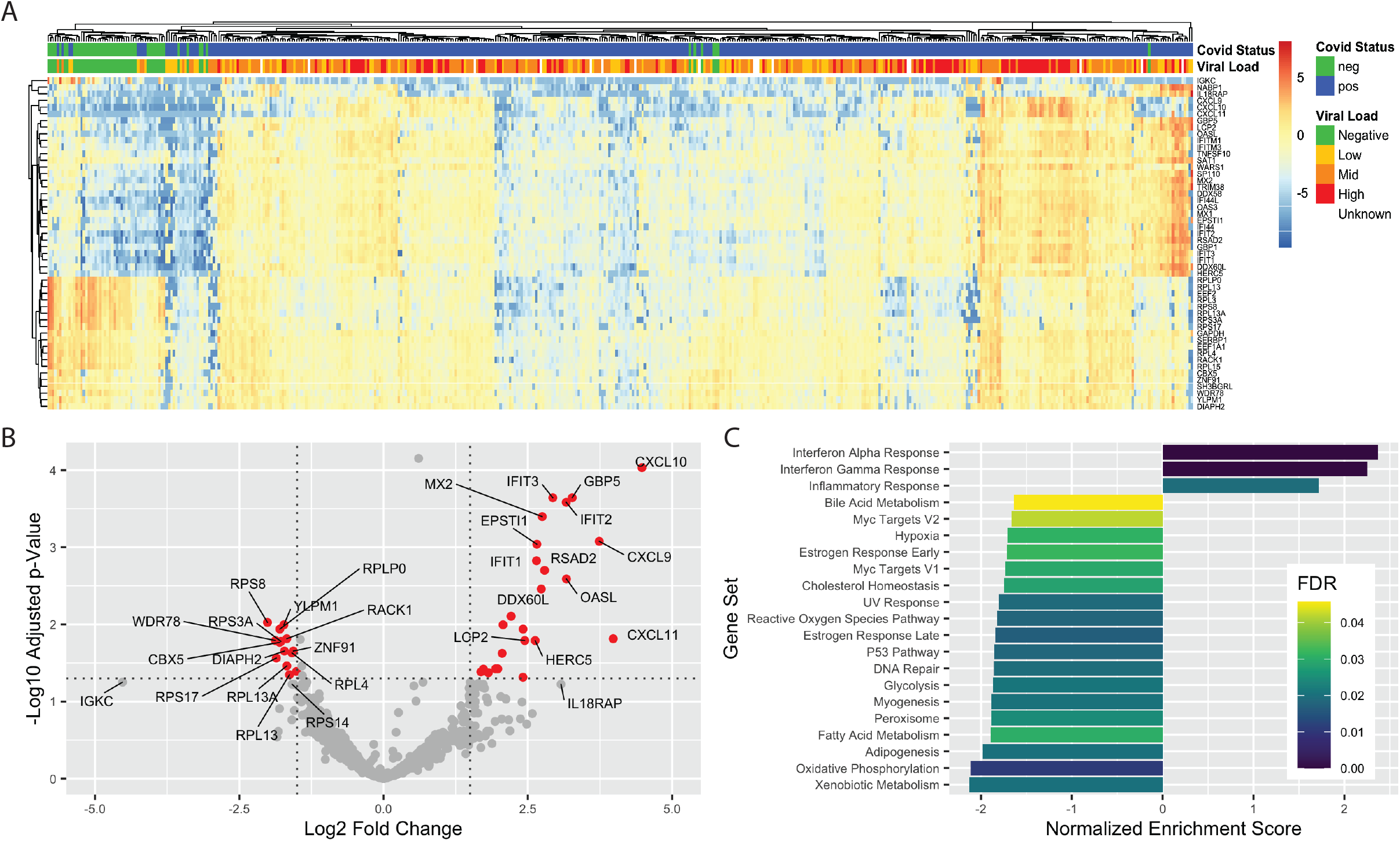
Differentially expressed genes in SARS-CoV-2 nasopharyngeal swabs. A) Clustering of samples based on 50 genes with the lowest adjusted pvalue. Log2 fold changes relative to gene mean are displayed by color. B) Volcano plot of 15 most upregulated and 15 most downregulated genes in SARS-CoV-2 positive samples relative to negative by log2 fold change. Red color indicates genes with log2 fold change > |1.5| and adjusted pvalue <0.05. C). Significant (FDR <0.05) pathways affected by SARS-CoV-2 infection identified by Gene Set Enrichment Analysis.

To interrogate the global regulatory and signaling programs induced by SARS-CoV-2 infection, we employed Gene Set Enrichment Analysis (GSEA) (21,22) of the 50 Hallmark Gene Sets of the Molecular Signatures Database (23). Sets with a significant (FDR < 0.05) positive enrichment score included Interferon Alpha, Interferon Gamma, and Inflammatory Responses (Figure 1C, Supplementary Figure 1A). Interestingly, we also found several metabolic pathways negatively enriched, including both Oxidative Phosphorylation and Glycolysis, suggesting a global reduction in production of proteins related to cellular energy production (Figure 1C, Supplementary Figure 1B). Broad downregulation of transcripts encoding metabolic machinery may represent either an antiviral response or viral-mediated disruption of host transcripts. We also performed a statistical enrichment test against the Biological Processes Gene Ontology (24,25). The most enriched processes (Supplementary Figure 1C-E) are related to either immune responses or translation. In addition to upregulation of innate antiviral transcripts (Figure 1B, Supplementary Figure 1D), we also found a consistent downregulation of transcripts encoding ribosomal proteins (Supplementary Figure 1E).

The SARS-CoV-2 receptor ACE2 is an interferon-regulated gene and is upregulated in response to SARS-CoV-2 infection (8). We examined the relationship between viral load, defined by the cycle threshold (Ct) of the N1 target during diagnostic PCR, and *ACE2*. We found that *ACE2* expression was associated with increased viral load: median counts of negative, low viral load (N1 ct > 24), medium viral load (N1 ct 24-19), and high viral load (N1 ct < 19) were 0, 1.93, 3.45, and 7.82, respectively (p = 7.46e^−13^, by Kruskal-Wallis one-way ANOVA; Figure 2A). A similar trend was found for other interferon-induced genes, a subset of which is shown in Figure 2A including those significantly upregulated in SARS-CoV-2 infection (*CXCL9*, *OASL*, *MX1*), negative regulators of inflammation (*CD274/PD-L1*, *USP18*), monocyte chemoattractant protein-1 (*CCL2*) (26). Conversely, the protease required for viral entry, *TMPRSS2*, was reduced upon viral infection but was not modulated by viral dose, nor were ribosomal proteins (*RPL4*, *RPS6*).

**Figure 2:**
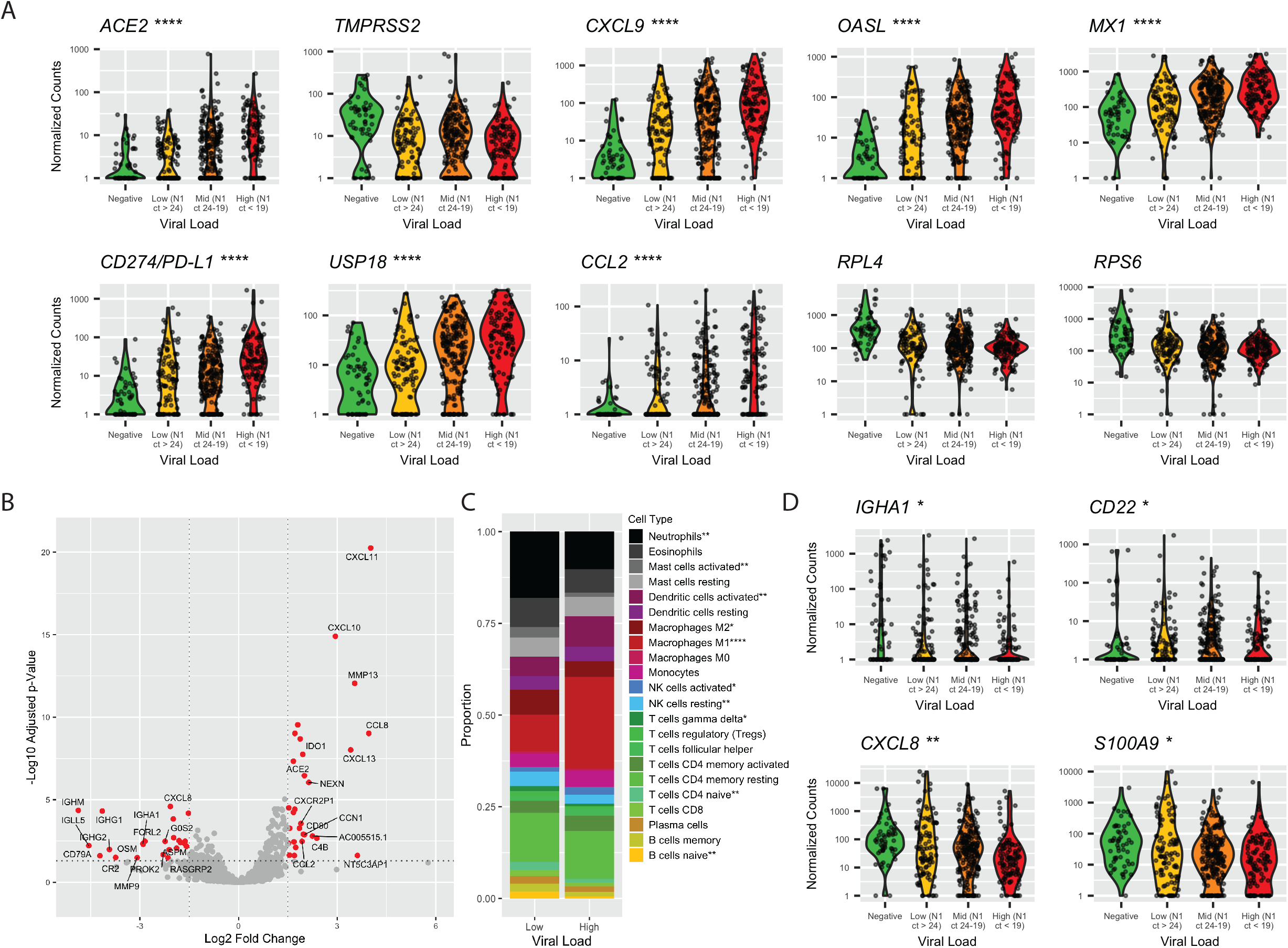
Differences in gene expression by SARS-CoV-2 viral load. A) Violin plots of select genes by viral load. Statistical significance between low and high viral load calculated by Mann Whitney U test, *p < 0.05, **p < 0.01, ***p < 0.001, ****p < 0.0001. B) Volcano plot of 15 most upregulated and 15 most downregulated genes in SARS-CoV-2 high viral load samples relative to low viral load by log2 fold change. Red color indicates genes with log2 fold change > |1.5| and adjusted pvalue <0.05. C) Proportion of cell types as a total of all immune cells, by CIBERSORTx. Significant differences in proportion of each cell type determined by T test, *p < 0.05, **p < 0.01, ***p < 0.001, ****p < 0.0001. D) Violin plots of B cell transcripts and neutrophil chemokine transcripts by viral load. Statistical significance between low and high viral load calculated by Mann Whitney U test, *p < 0.05, **p < 0.01, ***p < 0.001, ****p < 0.0001.

We next specifically examined gene expression differences between high (N1 ct < 19, n=108) relative to low (N1 ct > 24, n=99) viral load samples. Figure 2B highlights the 15 most upregulated and 15 most downregulated of the 363 total differentially expressed genes (adjusted pvalue < 0.1, Supplementary Table 2). While genes upregulated in high viral load samples were dominated by proinflammatory and/or interferon-induced factors such as *CXCL9/10*, *IDO1*, and *CD80*, genes with higher expression in low viral load samples included chemokines for neutrophils (*CXCL8, S100A9*), and B cell-specific transcripts (*FCRL2, IGHG1, IGHM, IGLL5, IGHG2, CD22*). Because this suggested differences in immune infiltration as a result of viral load, we performed in silico cell sorting of immune cells using CIBERSORTx (27) and found a higher proportion naïve B and T cells, neutrophils, and M2-polarized macrophages in low viral load samples (3.5, 2.2, 1.6, and 1.8 fold increased, respectively), while high viral load samples contained a larger proportion of M1 macrophages, activated NK cells, and activated dendritic cells (2.5, 1.6, and 1.6 fold upregulated, respectively; Figure 2C). Levels of transcripts encoding B cell proteins and neutrophil chemokines varied by viral load (Figure 2D).

Detection of differential infiltration of antigen-presenting cells and lymphocytes to the nasopharynx in high vs low viral load samples highlights the role immune cells play in the host response to SARS-CoV-2. To understand whether in vivo infection could be adequately modeled in vitro, we examined gene expression differences in human airway epithelial (HAE) cells 3 and 7 days post infection with SARS-CoV-2 and compared the DE genes at day 7 to those from SARS-CoV-2 positive vs negative (Figure 1) and high vs low viral load SARS-CoV-2 positive samples (Figure 2), resulting in a consensus set of 19 upregulated genes that define cell-intrinsic host antiviral responses to SARS-CoV-2 infection (Figure 3A). When this consensus set was tested for statistical enrichment in the DisGeNET (28) of disease ontologies, we found a high degree of overlap with influenza signature genes (Figure 3B), including a number of interferon-induced genes that mediate the acute antiviral response in the respiratory tract (Figure 3C). Notably, in the HAE cells, there was no sign of induction of an interferon response at 3 days post infection in spite of a 10-fold higher infectious dose of virus used and virus making up 0.3% of reads. At 7 days post infection, SARS-CoV-2 comprised 5.3% of reads.

**Figure 3:**
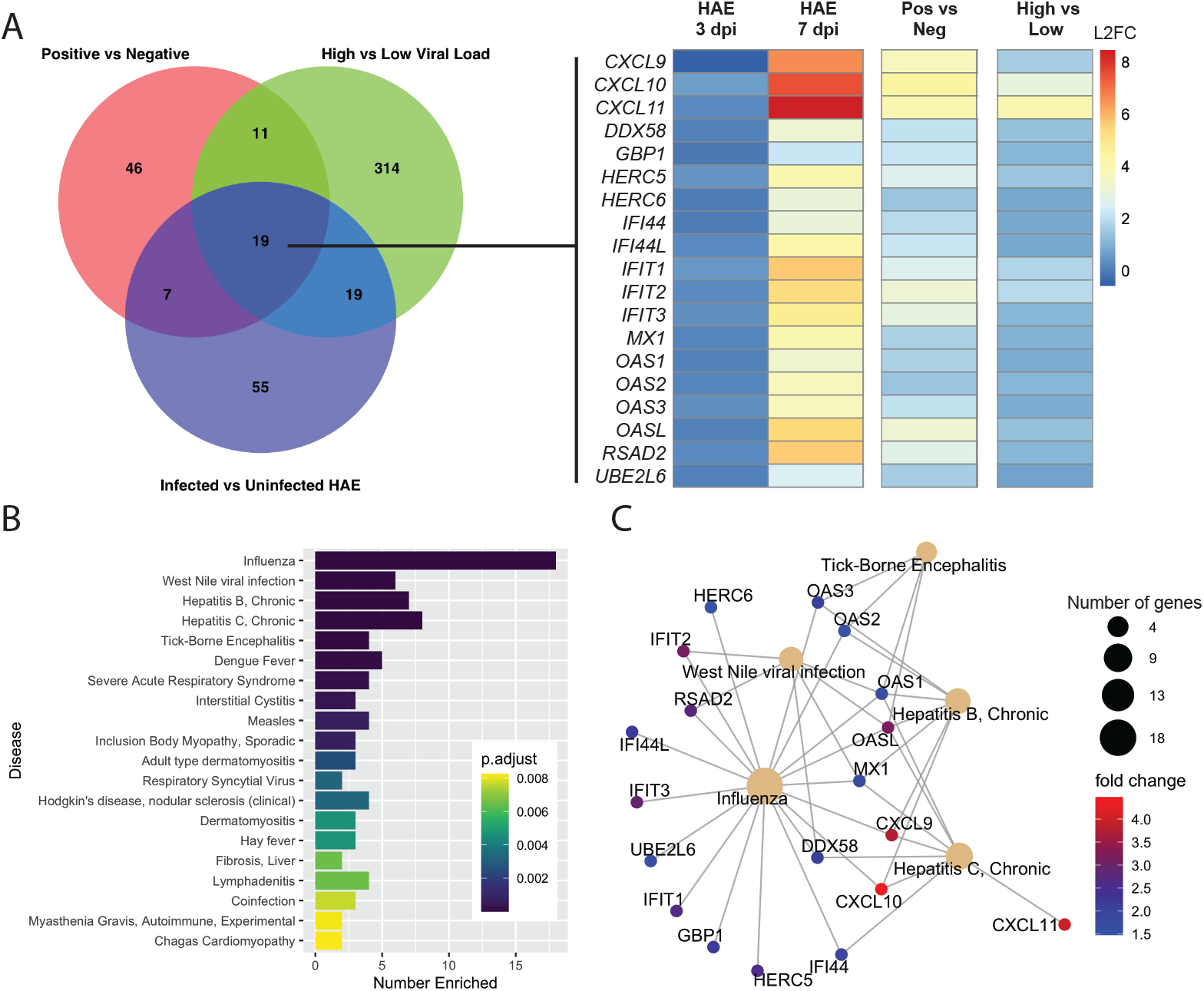
Consensus genes induced upon SARS-CoV-2 expression. A) Venn diagram of DE genes in SARS-CoV-2 positive vs negative, high vs low viral load, and top 100 genes with the highest absolute log2 fold change in infected vs uninfected HAE. Consensus set of 19 genes DE in all three analyses are shown, with log2 fold change values relative to uninfected HAE (for day 3 and day 7 post infection), SARS-CoV-2 negative NP swabs (for SARS-CoV-2 positive NP swabs), or low viral load (for high SARS-CoV-2 viral load samples). SARS-CoV-2 reads at day 3 and 7 post infection were 0.3% and 5.3%, respectively. B) Top 20 DisGeNET terms for which SARS-CoV-2 cell-intrinsic antiviral response consensus genes are overrepresented. Number Enriched is the number of SARS-CoV-2 consensus genes that belong to each disease term. C) Interaction network of SARS-CoV-2 consensus genes for top 5 most similar diseases identified in B. Size of disease node represents the number of genes enriched, and fold change is the log2 fold change seen in SARS-CoV-2 positive vs negative NP swabs.

Observed heterogeneity in host response to SARS-CoV-2 infection (Figure 1A) may be a result of co-infection or composition of the nasal flora. Proportion of reads assigned to virus, bacteria, or human in Supplemental Figure 2A shows a range of bacterial:human ratio among negative samples, while SARS-CoV-2 reads predominate at lower Ct values. Consistent with a dramatic reduction in respiratory virus transmission (29), presumably due to physical distancing measures enacted due to the SARS-CoV-2 pandemic, we found viral coinfections in only 14 of 430 SARS-CoV-2 positive samples (3.25%), and a single SARS-CoV-2 negative patient with a viral infection (2.5%) (Supplementary Figure 2B). We also found a number of samples enriched for potentially pathogenic bacterial components of the nasal flora, particularly *Moraxella catarrhalis* (RPM>100 in 3/37 (8.1%) SARS-CoV-2 negative, and 58/413 (14.0%) SARS-CoV-2 positive), although the clinical significance of this observation is uncertain. Even after SARS-CoV-2 reads were subtracted from each sample, we found that high viral load samples had a significantly lower burden of bacteria than mid or low viral load, or SARS-CoV-2 negative samples (Mann Whitney p = 0.0014, 0.0067, 0.00028, respectively; Supplementary Figure 2C).

Infection time course may account for the observed differences in immune related genes in high vs low viral load samples: Patients receiving repeat SARS-CoV-2 testing have a reduced viral load over time (13,30,31). Although we do not have the ability to tie our data from our large set of positive samples back to the onset of symptoms, we have seen gene expression changes in a small set (n=3) of matched longitudinal samples with a mean elapsed time of 6.3 days between collections and a mean increase in Ct value of 5.29, representing a 39-fold reduction in viral load (Figure 4A). Enriched GO terms include those related to translation and immune regulation (Figure 4B). Notably, in the second sample collected, we saw increases in genes such as *C1QA, -B*, and -*C* and *HLA-DQB1* that drive humoral immune responses and those involved in wound healing (*APOE, CD36, RHOC*), as well as reductions in negative regulators of each process (ie, *TREM1, TFPI*). We also tested the 19 SARS-CoV-2 signature genes (Figure 3A), and found reductions in most at the second collection timepoint, although only *RSAD2*, *IFIT2*, and *HERC5* decreases were statistically significant with three samples. We also saw a recovery in the expression of ribosomal proteins over time (Figure 4C). Analysis of data from additional patients with more extensive

**Figure 4:**
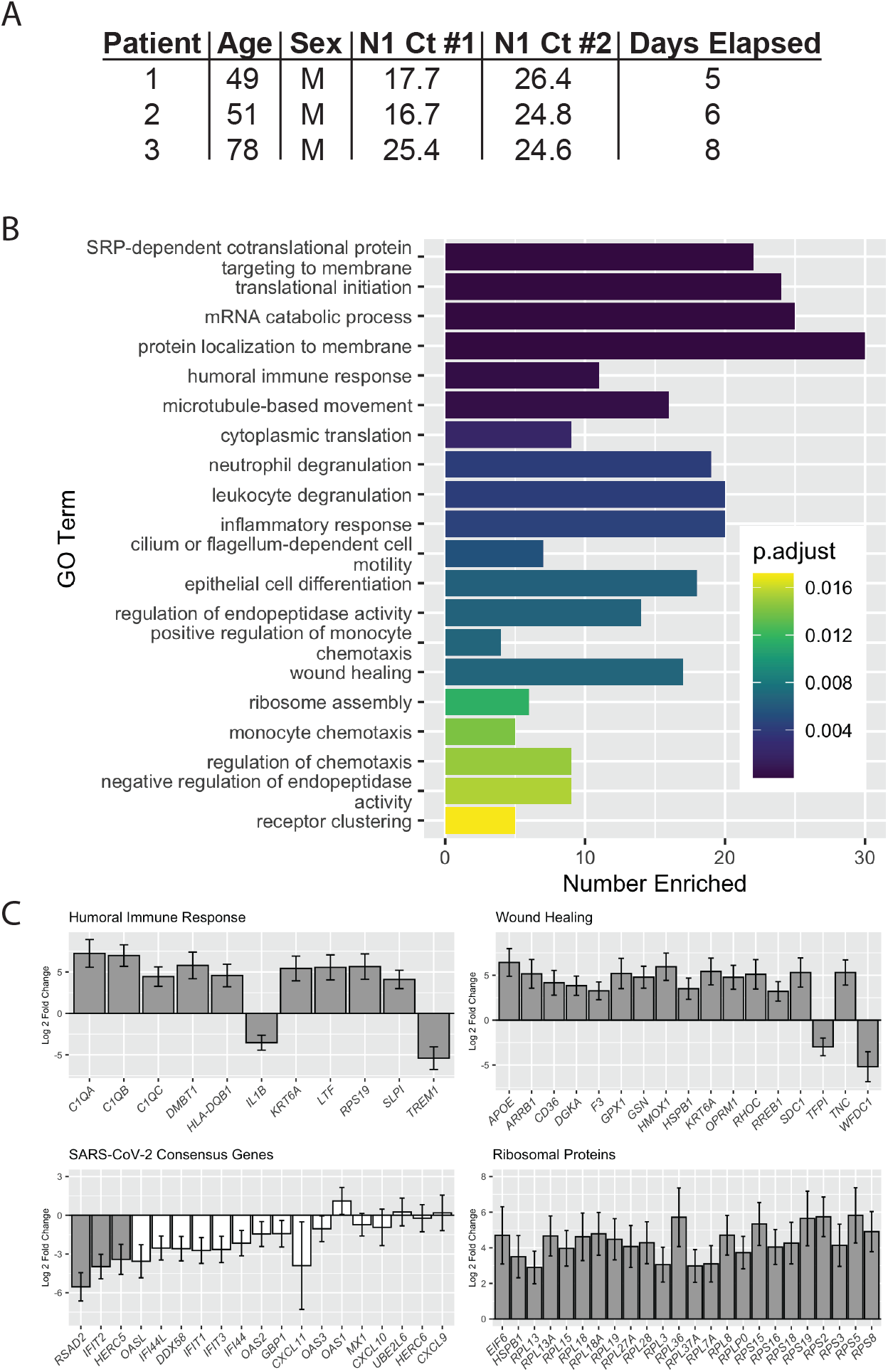
Differentially expressed genes in patient-matched longitudinal samples. A) Patient demographics information for longitudinal samples. B) Top 20 Biological Process Gene Ontology terms for which longitudinal DE genes are overrepresented. Number Enriched is the number of DE genes that belong to each GO Term. C) Log2 fold changes for DE genes in Humoral Immune Response and Wound Healing GO Terms, consensus antiviral SARS-CoV-2 genes, and ribosomal proteins. Grey bars: padj < 0.1, white bars: padj > 0.1.

Clinically, COVID-19 cases tend to be more severe for older adults and males (1). No significant difference in N1 Ct was observed based on age or sex (Figure 5A). To understand the differences in host response to SARS-CoV-2 infection, we tested the interaction between infection and age (greater than 60), controlling for non-infection-related age differences with SARS-CoV-2 negative samples. We found only two genes altered as a result of the interaction between age and SARS-CoV-2 infection: a 30-fold reduction in production of CXCL11 (Figure 5B), an interferon-induced chemokine for natural killer and CD8+ T cells, and a 17-fold reduction in polycomb group factor 6 (*PCGF6*) (Supplementary Figure 3A), a polycomb repressor complex protein known to play a role in repression of dendritic cell activation (32). Although we did not find additional genes altered specifically as a result of age in SARS-CoV-2 infection, we did find that *CXCL9* and C*XCL10* are not induced as strongly in SARS-CoV-2 positive patients age 60 or higher. We also found reduced expression of the receptor for *CXCL9/10/11, CXCR3*, the apoptosis-inducing factor *GZMB* secreted by NK and T cells, and the effector T cell marker, *CD8A*. This data suggests that age-related T and NK cell dysfunction (33,34) may play a role in SARS-CoV-2 pathogenesis in older individuals.

**Figure 5:**
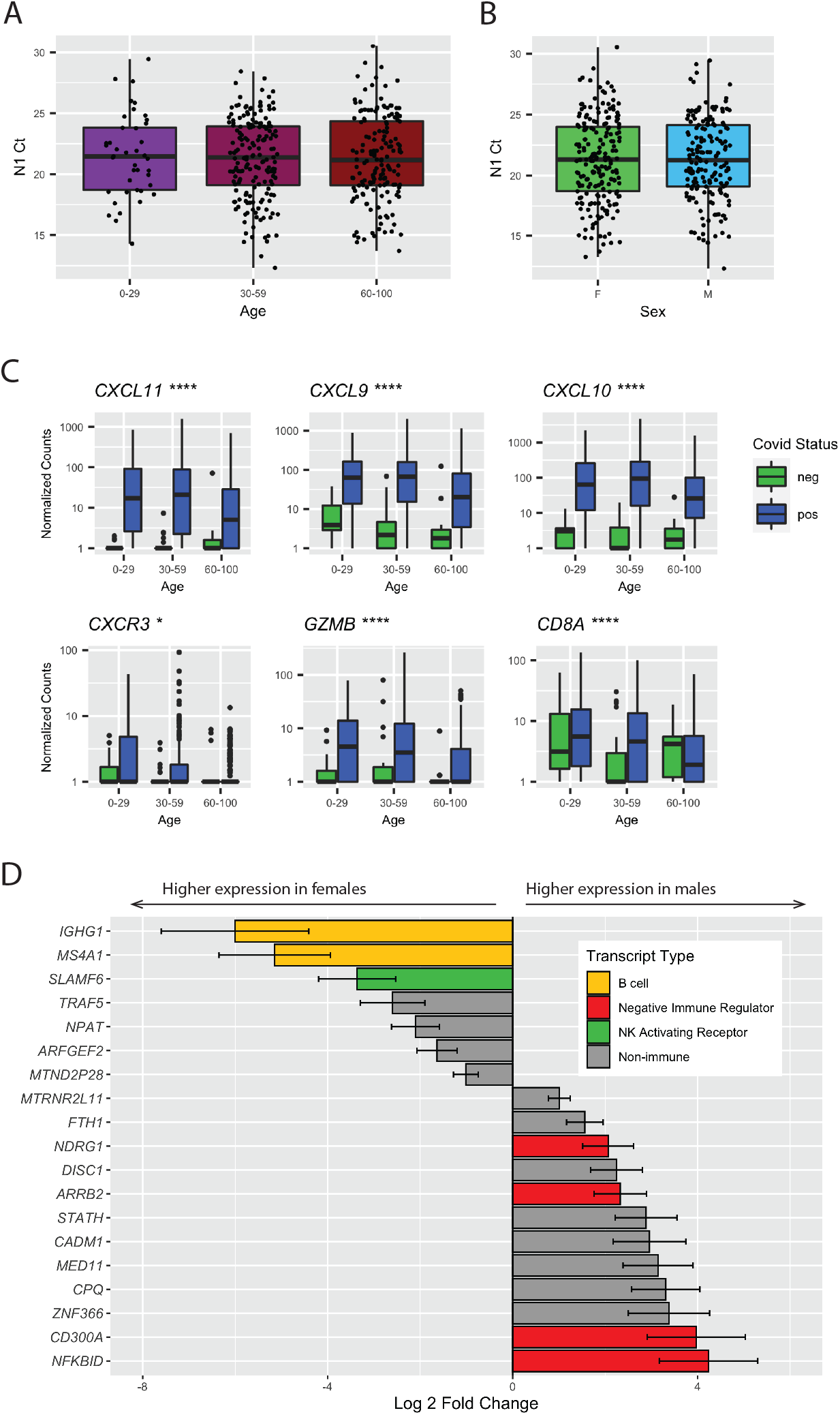
Age and sex cause differences in gene expression upon SARS-CoV-2 infection. A) N1 Ct values by age group. No significant differences between were observed by Kruskal-Wallis ANOVA. B) N1 Ct values by sex. No significant difference between groups was observed by t-test. C) Gene expression differences by age and viral load. Significance by Mann Whitney U test between SARS-CoV-2 positive samples age >60 and <60 is shown, *p < 0.05, **p < 0.01, ***p < 0.001, ****p < 0.0001. D) Sex-modulated DE genes (padj <0.1) upon SARS-CoV-2 infection. Genes elevated in females are shown as negative log2 fold changes, and those elevated in males as positive log2 fold changes.

We performed a similar analysis to evaluate sex differences in SARS-CoV-2 infection and found a total of 19 genes for which the differences in expression based on sex could be attributed to SARS-CoV-2 infection. Supplementary Figure 3B highlights the top 10 non-redundant enriched GO categories, most of which are related to immune function. In men, we found downregulation of B cell-specific transcripts (*IGHG1* and *MSA4A/CD20*), downregulation of the NK-activating receptor *SLAMF6*, and an upregulation of several inhibitors of NFκB signaling (*NDRG1, ARRB2, CD300A*, and *NFKBID*) (Figure 5D). The downregulation of B cell-specific markers suggests differences in lymphocyte composition and/or trafficking in males. Furthermore, the reduction in NK cell activating receptors and upregulation of negative regulators of immune effector function, and resultant throttling of effector function, is consistent with a more severe manifestation of COVID-19 in males.

## Discussion

One of the hallmarks of COVID-19 is a dysregulated antiviral immune response. Studies of SARS-CoV, which also employs *ACE2/TMPRSS2*-mediated entry, have demonstrated infection does not always result in production of interferon-β in macrophages and dendritic cell (35), and significantly delayed expression of type II or III interferon in lung cells (10). Moreover, infection of BALB/c mice with SARS-CoV did not result in detectable IFNβ until 24 hours, at which point viral titers had nearly reached a peak; lung damage resulting from the subsequent massive infiltration of inflammatory macrophages could be abrogated by pre-treatment with type I interferons (11). Similar viral kinetics have been observed in SARS-CoV-2-infected patients (13,14), and ferrets (12). Collectively, these results support a common mechanism by which SARS-CoV and SARS-CoV-2 suppress intracellular viral detection and subsequent interferon induction long enough for viral replication to occur.

Our transcriptomic analysis of nasopharyngeal swabs reveals the robust induction of an interferon response by SARS-CoV-2 infection (Figure 1), similar to that observed by Butler et al (20). The highest levels of individual interferon-responsive genes were seen in samples with the highest viral load (Figure 2A) and enriched for transcripts associated with inflammatory macrophages and activated DCs and NK cells, three primary sources of type I and II interferons (Figure 2B). When repeat swabs were taken from patients with an average 6.3 day time period between sampling, the interferon response had waned, as had viral load (Figure 4C). Notably, in contrast to the robust expression of interferon-regulated transcripts in HAE seven days after infection with SARS-CoV-2, there was limited evidence of induction of an interferon response after only three days, consistent with a SARS-CoV-like functional repression of interferon signaling.

COVID-19 patients frequently develop interleukin-6-driven cytokine release syndrome (CRS), and elevated serum IL-6 correlates with respiratory failure and poor clinical outcomes (36). Treatment with the IL-6 receptor blocking antibody tocilizumab has effectively treated COVID-19 symptoms in some patients (37). We did not see a significant difference in expression of *IL-6*, nor of other CRS-associated factors such as *TNF* or *VEGF*, when we analyzed SARS-CoV-2 positive samples relative to negative, nor in high vs low viral load SARS-CoV-2 positive samples. This could be attributed to the nasopharynx not being a particularly sensitive anatomic location to probe markers of systemic inflammation compared to serum or lower respiratory sites. Additionally, our choice to use a large number of samples at relatively low sequencing depth likely reduced our sensitivity to detect differences in low-abundance and short-lived transcripts like cytokines.

One of the more striking patterns we observed is the marked downregulation of transcription of ribosomal proteins upon SARS-CoV-2 infection (Figure 1B), and the recovery of expression during disease progression (Figure 4C). Global inhibition of host transcription is a strategy employed by many viruses via diverse mechanisms such as disrupting transcriptional pre-initiation complex assembly (38,39) or cleavage of TATA-binding protein (40). MERS-CoV and SARS-CoV nsp1 both cause decay of host mRNA (41,42); in MERS-CoV, host mRNA degradation results from an endonucleolytic function of nsp1 itself (43). Nsp1 from SARS-CoV and SARS-CoV-2 share 84% amino acid identity, therefore it is likely that SARS-CoV-2 nsp1 can also function directly or indirectly to promote host RNA degradation. Global downregulation of host transcription may also be driven in part by SARS-CoV-2 ORF6 protein, which binds to the mRNA export factor RAE1 and nuclear pore protein Nup98 in a similar manner as the VSV M protein (44), or the ORF7a protein, which binds to proteins involved in ribosomal assembly and nuclear export (44).

Metagenomic analysis of SARS-CoV-2 positive samples revealed a low rate of viral co-infection (3.25%), consistent with 3.2% reported by Butler et al in New York City (20). This is likely due to a dramatic reduction in circulating respiratory viruses in March and April 2020 caused by physical distancing measures across the country. Among the samples from which we could obtain high quality metagenomic data, we found 46 of 413 SARS-CoV positive samples but only 1 of 37 SARS CoV negative samples extensively colonized (RPM>1000) by potentially pathogenic bacteria (Supplementary Figure 2B), although no pattern of infection was found based on viral load. More work is required to understand how these normalized read count thresholds for resident nasopharyngeal microbiota correlate with transitions from colonization to mild illness to potentially invasive disease.

Finally, understanding age- and sex-related differences in responses to SARS-CoV-2 infection is of critical importance as approximately 90% of SARS-CoV-2 deaths in Washington State have been seen in individuals over 60. Our data show that in individuals over 60, expression of interferon-induced chemokines is reduced, possibly contributing to a reduction in transcripts for cytotoxic T and NK cells. Immune dysfunctions in older individuals are well-characterized (2,3,33,34), and likely contribute to poorer COVID-19 outcomes; results from clinical trials of type I and III interferons in severely ill patients are likely to further define the role of interferon signaling in older adults (45–47).

Differences in immune responses in males and females are due to a variety of factors, including the effects of sex hormones and the X-linked nature of many immune genes (48). The bias towards expression of B cell transcripts in females in our study is consistent with higher levels of B cells in females regardless of age (49). Females also tend to have increased inflammation in response to viral infections (4). The observed increased expression of inhibitors of NFkB in males with SARS-CoV-2 may represent either inappropriate throttling of the antiviral immune response or an adaptive mechanism to reduce deleterious inflammation, a hallmark of COVID-19 pathogenesis.

Collectively, we demonstrate induction of an antiviral response characterized by type I and II interferon induction, which wanes with time and is correlated with viral load. We also find evidence of transcriptional repression by SARS-CoV-2. Lastly, we show that differences in immune responses may underlie disparities in outcomes for two higher risk groups, males and the elderly.

## Methods

### IRB Approval

Sequencing of excess clinical samples was approved by the University of Washington IRB (STUDY00000408).

### Sample Collection, RNA extraction, and qPCR

NP swabs of patients with suspected SARS-CoV-2 infection were collected in 3 mL viral transport medium (VTM). Total RNA was extracted from 200 or 140 μL of VTM using either the Roche MagNAPure or Qiagen BioRobot automated platforms, respectively (50). Quantitative PCR for the SARS-CoV2 N1 target was performed on the Applied Biosystems 7500 real time PCR instrument (51,52).

### Library preparation and sequencing

Metagenomic next-generation sequencing (mNGS) was performed as previously described (17,53). Briefly, 18 μL of extracted RNA was treated with Turbo DNAse (ThermoFisher). First strand cDNA synthesis was completed using SuperScript IV (ThermoFisher) and random hexamers (Invitrogen) followed by second strand synthesis by Sequenase V2.0 (ThermoFisher). The resulting cDNA was purified using either the DNA Clean & Concentrator kit (Zymo) or 1.6x volumes of AMPure XP beads (Beckman Coulter). Library preparation was performed using the Nextera XT Kit (Illumina). Libraries were cleaned with 0.7x or 0.75x volumes of Ampure beads (Beckman Coutler), quantified using either the Qubit dsDNA HS assay (ThermoFisher) or Quant-iT dsDNA HS assay (ThermoFisher), quality checked by Bioanalyzer or TapeStation (Agilent), pooled, and sequenced on 1 × 75 bp runs on an Illumina NextSeq or 1 × 101 bp runs on an Illumina NovaSeq.

### Pseudoalignment

Raw FASTQ files were adapter and quality trimmed by Trimmomatic v0.39 (54) using the call “leading 3 trailing 3 slidingwindow:4:15 minlen 20”. Trimmed reads were pseudoaligned to the Ensembl v96 human transcriptome using Kallisto v0.46 (55) assuming an average library size of 300+/−100 base pairs. Only samples with more than 500,000 pseudoaligned reads were used for RNAseq analysis.

### Differential Expression

Pseudoaligned reads were pre-filtered to remove any genes with average expression of less than one read per sample, then normalized and differential expression calculated with the R package DEseq2 v1.28.1 (56). Correction for batch effects was incorporated into the design formula and modeling performed using the Wald test with outlier replacement. Results were deemed significant at a Benjamini-Hochberg adjusted pvalue <0.1. Gene expression differences attributable to sex or age were incorporated into the design formula as interaction terms.

### Gene Set Enrichment Analysis (GSEA)

GSEA was performed on normalized counts on GSEA Software version 4.0.3 (21,22). Gene ranking was generated with the Signal2Noise metric and analyzed against the mSigDB Hallmarks v7.1 gene sets (23).

### Metagenomics

Metagenomic analysis of the RNA sequence was performed using CLOMP v0.1.4 (17) with the default options and visualized using the Pavian metagenomic explorer (57). Viral species level taxonomical classifications with an RPM greater than 25 were confirmed via BLAST v2.10.1 (e-value 1e-5).

#### Human Airway Epithelial (HAE) cultures

The EpiAirway AIR-100 system (MatTek Corporation) consists of normal human-derived tracheo/bronchial epithelial cells that have been cultured to form a pseudostratified, highly differentiated mucociliary epithelium closely resembling that of epithelial tissue in vivo. Upon receipt from the manufacturer, HAE cultures were transferred to 6-well plates containing 1.0 ml EpiAirway medium per well (basolateral feeding, with the apical surface remaining exposed to air) and acclimated at 37°C in 5% CO2 for 24 hours prior to experimentation.

#### Viral growth in HAE

HAE cultures were infected by applying 200 μl of EpiAirway phosphate-buffered saline (MatTeK TEER Buffer) containing 2,000 PFU or 20,000 PFU of infectious clone-derived SARS-CoV-2 expressing a stable mNeonGreen reporter gene (icSARS-CoV-2-mNG) (58) to the apical surface for 90 min at 37°C. At 90 min, the medium containing the inoculum was removed, the apical surface was washed with 200 μl of TEER buffer, and cultures were placed at 37°C. Cultures were fed every other day with 1.0 ml medium via the basolateral side. Media was removed, and cultures were lysed with TRIzol Reagent (ThermoFisher) at three days post infection (20,000 PFU challenge) and at 7 days post infection (2,000 PFU challenge). Bam files of viral sequence are deposited in the sequence read archive, NCBI Bioproject PRJNA634194.

### HAE RNAseq and analysis

RNA from uninfected and infected HAE was extracted using Direct-zol RNA MicroPrep (Zymo). Libraries were generated using the TruSeq Stranded mRNA kit (Illumina) and 2×100bp paired-end reads sequenced on a Novaseq. Pseudoalignment using Kallisto v0.44 and differential expression analysis was performed as above.

### Statistics and visualization

All calculations were performed in R v4.0.0. Statistical enrichment tests of Gene Ontology (24,25) and DisGeNET (28) pathways were performed in the clusterProfiler R package (59). Images were generated using packages including DOSE (60), ggplot2, pheatmap, and VennDiagram.

## Data Availability

Raw counts and metadata for each nasopharyngeal sample is deposited in the NCBI Gene Expression Omnibus GSE152075.

## Acknowledgements

The authors thank Amin Addetia and Joshua Lieberman for their thoughtful comments to improve the manuscript. This work was supported by funding from the National Institutes of Health (AI146980, AI121349, and NS091263 to MP) and the Department of Laboratory Medicine at the University of Washington School of Medicine.

---

**Supplementary Figure 1:**
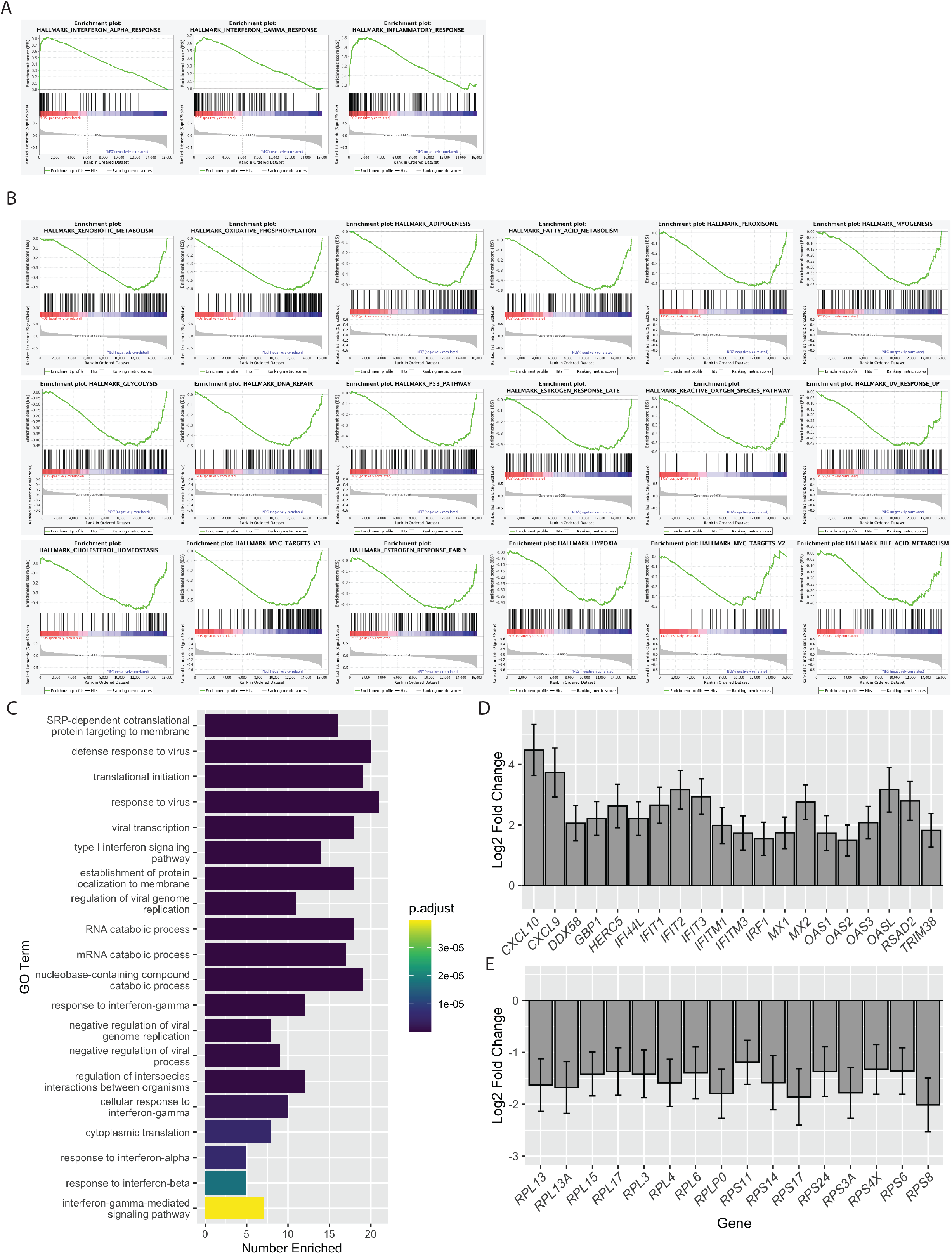
Differentially expressed Gene Sets and Gene Ontology Biological Process Terms in SARS-CoV-2 nasopharyngeal swabs. A) Enrichment plots of gene sets significantly (FDR<0.05) positively enriched in SARS-CoV-2 samples. B) Enrichment plots of gene sets significantly (FDR<0.05) negatively enriched in SARS-CoV-2 samples. C) Top 20 Biological Process Gene Ontology terms for which differentially expressed genes in SARS-CoV-2 samples are overrepresented. Number Enriched is the number of SARS-CoV-2 differentially expressed genes that belong to each GO Term D) Fold change of genes belonging to GO Term “defense response to virus”. E) Fold change of genes belonging to GO Term “SRP-dependent cotranslational protein targeting to membrane”.

**Supplementary Figure 2:**
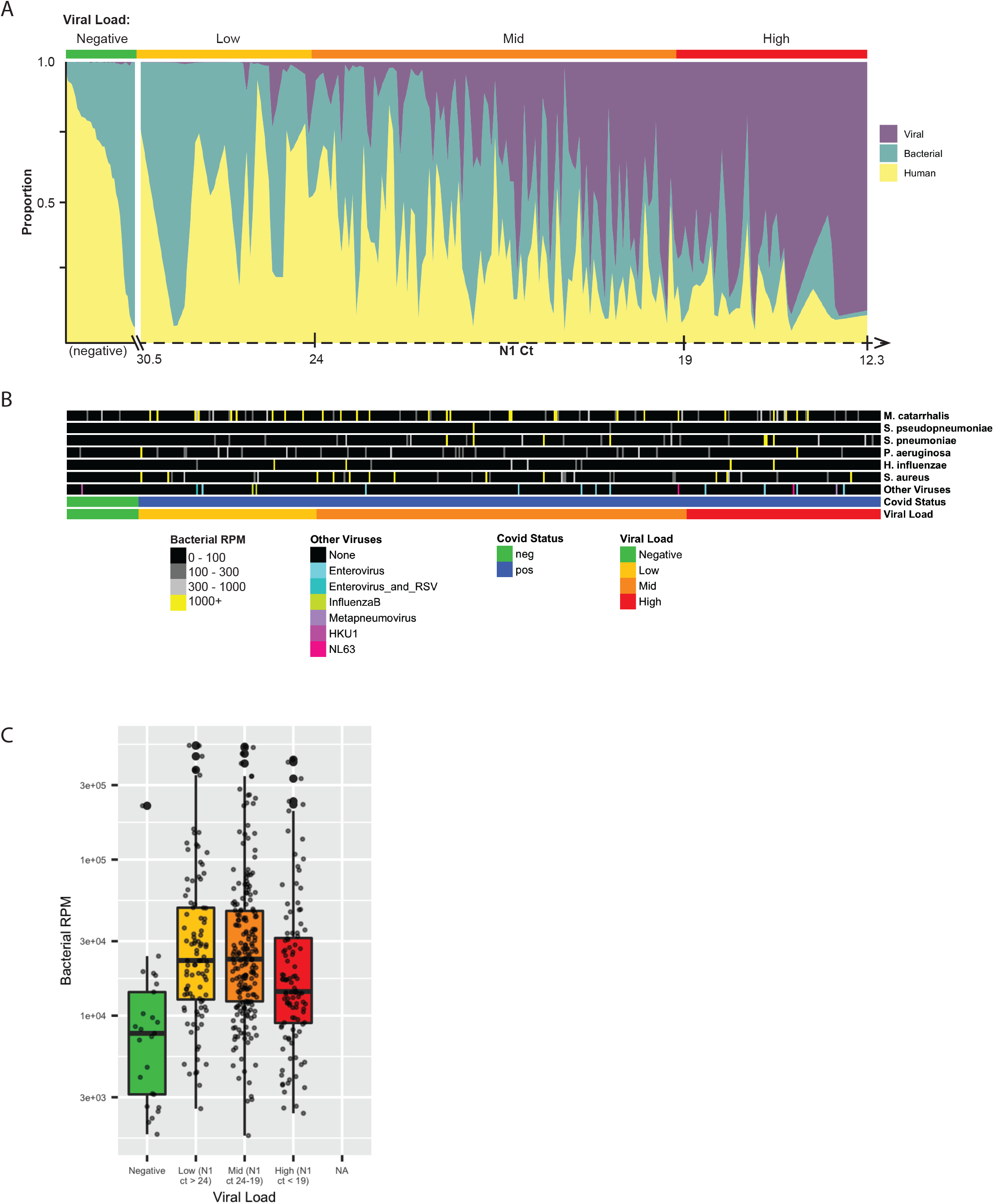
Metagenomic analyses of SARS-CoV-2 positive and negative samples. A) Loess-smoothed area plot showing the proportion of human, viral, and bacterial reads for each sample. Positive samples are arranged in reverse order of N1 Ct. B) Colonization and co-infection of non-SARS-CoV-2 respiratory viruses and clinically relevant bacterial species. C) Violin plot of bacterial RPM after correcting by subtraction of SARS-CoV-2 reads.

**Supplementary Figure 3:**
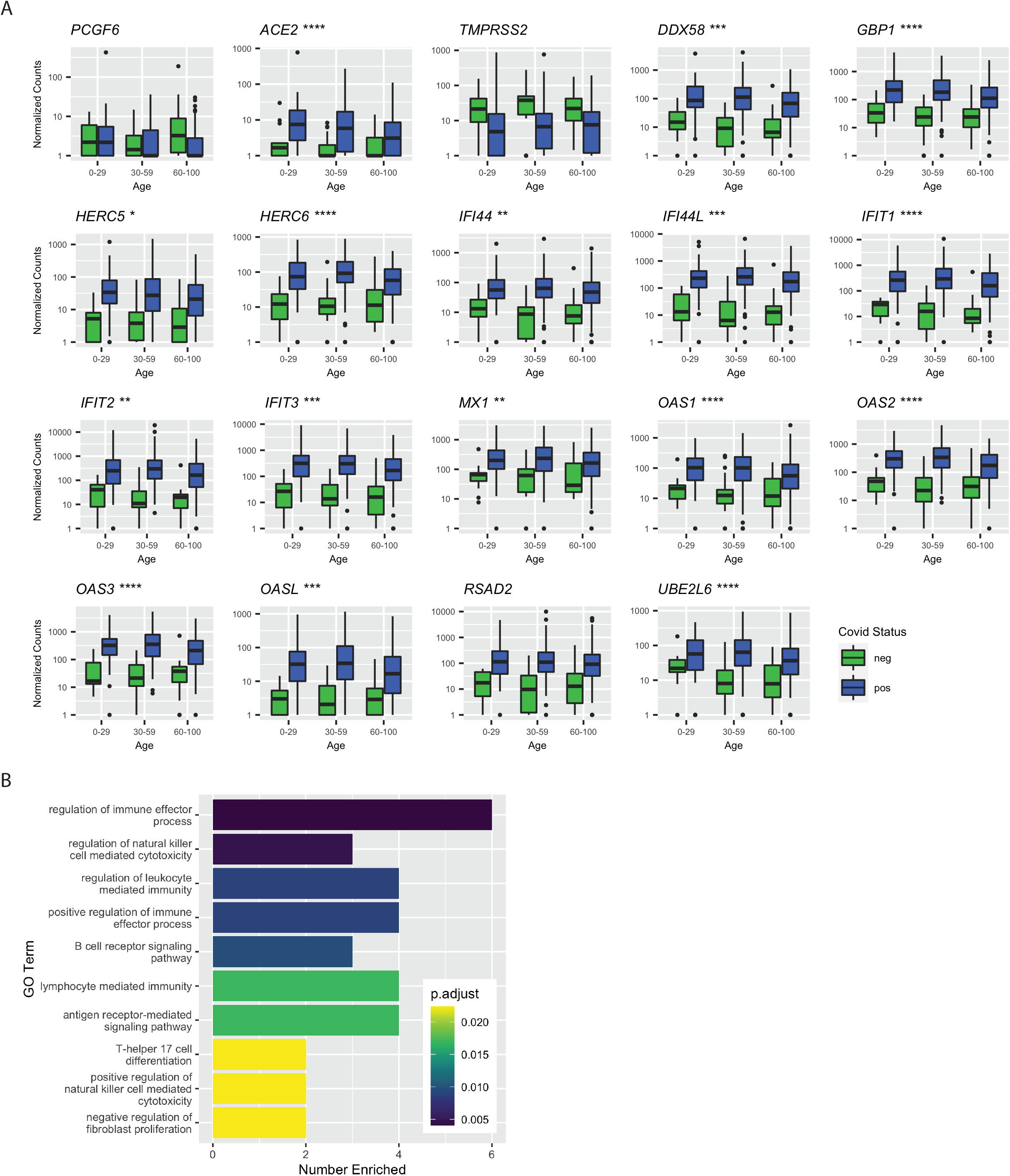
Age and sex differences in gene expression upon SARS-CoV-2 expression. A) Gene expression differences by age and viral load. Significance by Mann Whitney U test between SARS-CoV-2 positive samples age >60 and <60 is shown, *p < 0.05, **p < 0.01, ***p < 0.001, ****p < 0.0001. B) Top 10 Biological Process Gene Ontology terms in which genes defining the male vs female response to virus are overrepresented.

**Supplementary Table 1**: Differentially expressed genes in SARS-CoV-2 positive samples relative to negative.

**Supplementary Table 2**: Differentially expressed genes in SARS-CoV-2 high viral load samples relative to low viral load.

